# Doomed by popularity: The broad use of the *Pm8* resistance gene in wheat resulted in hypermutation of the *AvrPm8* gene in the powdery mildew pathogen

**DOI:** 10.1101/2022.08.24.505094

**Authors:** Lukas Kunz, Alexandros G. Sotiropoulos, Johannes Graf, Mohammad Razavi, Marion C. Müller, Beat Keller

## Abstract

- The *Pm8* resistance gene against the powdery mildew disease has been introgressed from rye into wheat as part of a large 1BL.1RS chromosomal translocation. Due to its high agronomic value, this translocation has seen continuous global use since the 1960’s on large growth areas, even after *Pm8* resistance was overcome. This allows studying the effect of long and widespread resistance gene use on a pathogen population.
- Using genome wide association studies in a global population of wheat mildew isolates, we identified the avirulence effector *AvrPm8* specifically recognized by *Pm8*.
- Haplovariant mining in the global population revealed 17 virulent haplotypes of the *AvrPm8* gene that grouped into two categories. The first one from geographically diverse regions comprised two single amino acid polymorphisms at the same position in the AvrPm8 protein, which we confirmed to be crucial for recognition by Pm8. The second category consisted of numerous destructive mutations to the *AvrPm8* open reading frame.
- Most individual gain-of-virulence mutations were found in geographically restricted regions, indicating they occurred recently as a consequence of the frequent *Pm8* use. We conclude that both standing genetic variation as well as locally occurring new mutations contributed to the global breakdown of the *Pm8* resistance gene introgression.

## Introduction

Wheat is one of the most widely cultivated crop species worldwide, serving as an important source of calories and protein for human nutrition (FAO, 2021). Sustainable wheat production is however threatened by numerous, fast evolving fungal pathogens. Breeding efforts continuously aim to incorporate new resistance traits into high yielding cultivars. Since the beginning of the twentieth century, introgressions from closely related wild and domesticated grass species represent one of the most valuable sources for new resistance genes (Wulff & Moscou, 2014). In the 1930’s, the 1RS chromosomal segment of rye (*Secale cereale*) cultivar ‘Petkus’ was introduced into hexaploid wheat, replacing the wheat chromosome arm 1BS (Lein, 1975). Cultivars carrying the 1BL.1RS translocation not only exhibited higher yield potential but also increased disease resistance against leaf rust, yellow rust, stem rust and wheat powdery mildew since the rye translocation harbors the *Lr26*, *Yr9*, *Sr31* and *Pm8* resistance genes (Crespo-Herrera *et al*., 2017). Due to the unique combination of favorable traits, cultivars carrying the 1BL.1RS translocation have been broadly used in wheat growing areas worldwide since the 1960’s and continue to be dominantly represented in wheat breeding programs in many countries (Lukaszewski, 1990; Rabinovich, 1998; Graybosch, 2001; Zhou *et al*., 2007). For example, in 1998 the 1BL.1RS translocation was present in 50% of the high yielding bread wheat lines from the ‘International Maize and Wheat Improvement Center (CIMMYT)’, while it reached up to 90% of planted hectarage in some national breeding programs (Rabinovich, 1998; Villareal *et al*., 1998; Purnhauser *et al*., 2011). While the *Sr31* gene remained effective against stem rust for more than 30 years before it was overcome by the highly virulent strain Ug99 (Pretorius *et al*., 2000; Singh *et al*., 2011), the mildew resistance gene *Pm8* broke down quickly in many regions of the world, usually within few years of large-scale deployment of 1BL.1RS cultivars (Bennett, 1984; Streckeisen, 1985; Namuco *et al*., 1987; Heun & Friebe, 1990; Purnhauser *et al*., 2011). Despite the quick reduction in resistance effectiveness, *Pm8* remained common within the wheat breeding pool, due to complete genetic linkage with the other favorable traits present on the 1BL.1RS translocation (Rabinovich, 1998; Purnhauser *et al*., 2011).

The *Pm8* gene is allelic to *Pm17*, a second rye-derived resistance gene used in wheat breeding and residing on a 1AL.1RS translocation from rye cultivar ‘Insave’ (Singh *et al*., 2018). *Pm8* and *Pm17* encode for intracellular, nucleotide-binding leucine-rich repeat (NLR) immune receptors and were found to be homologous to the endogenous wheat *Pm3* resistance locus (Hurni *et al*., 2013; Singh *et al*., 2018). The *Pm3* resistance gene encodes for numerous, highly similar NLR variants (*Pm3a* to *Pm3t*) (Yahiaoui *et al*., 2004; Bhullar *et al*., 2009) that confer resistance against wheat powdery mildew *Blumeria graminis* f. sp. *tritici* (*B.g. tritici*). *Blumeria graminis* (grass powdery mildew) is an obligate biotrophic ascomycete fungus existing in numerous sublineages (formae speciales) that exhibit high levels of host specificity, such as *B.g. tritici* exclusively infecting wheat or *B.g. secalis* growing on the *Pm8/Pm17* donor species rye. The NLR proteins encoded by the *Pm3* allelic series and *Pm17*, where shown to provide race specific resistance against *B.g. tritici* through recognition of mildew encoded effector proteins (avirulence factors, *AVRs*) (Brunner *et al*., 2010; Bourras *et al*., 2015; Bourras *et al*., 2019; Müller *et al*., 2022). The *B.g. tritici* avirulence genes *AvrPm3^a2/f2^*, *AvrPm3^b2/c2^*, *AvrPm3^d3^* and *AvrPm17*, recognized by *Pm3a/Pm3f*, *Pm3b/Pm3c*, *Pm3d* and *Pm17*, respectively encode for highly diverse, small, secreted effector proteins with a common Y/FxC motif and a predicted RNA-se like structure. Population level sequence analysis coupled with functional characterization of gain-of-virulence variants of these *AVRs* has provided a detailed insight into the evolutionary and molecular mechanisms involved in the resistance breakdown of *Pm3* and *Pm17*. For example, gain-of-virulence mutations in the *AvrPm3* and *AvrPm17* effector genes were exclusively found to generate single amino acid polymorphisms, likely allowing *B.g. tritici* to evade NLR recognition while preserving effector virulence function (McNally *et al*., 2018; Bourras *et al*., 2019; Müller *et al*., 2022).

It is estimated that about 5% of modern bread wheat lines harbor a *Pm3* resistance gene (Bhullar *et al*., 2010) and *Pm17* was mostly used in limited geographic regions such as the US or China (Zeng *et al*., 2014; Graybosch *et al*., 2019). Thus, *Pm8* is by far the most frequently and continuously used mildew resistance gene in wheat breeding and agricultural production of the last decades (Rabinovich, 1998; Villareal *et al*., 1998; Purnhauser *et al*., 2011; Hurni *et al*., 2013). The identification of the corresponding avirulence gene *AvrPm8* is therefore of high relevance, as it would provide an unprecedented insight into the consequences of such broad resistance gene application on the global wheat mildew population. Simultaneously it would allow to better understand the rapid breakdown of *Pm8* resistance that occurred within few years of its deployment and seemingly independently in multiple regions of the world. Such insights would guide more-informed decisions in future resistance breeding, resulting in more durable resistance against the powdery mildew pathogen.

## Results

Genome wide association studies (GWAS) have been previously used to identify avirulence genes in *B.g. tritici* (Praz *et al*., 2017; Bourras *et al*., 2019). In order to identify the *AvrPm8* gene we performed GWAS, using a diversity panel of 79 *B.g. tritici* isolates from a worldwide collection (Sotiropoulos *et al*., 2022) and the chromosome-scale genome assembly of *Pm8* avirulent isolate ISR_7 (Müller *et al*., 2022) as a reference (Fig. 1a). Since *Pm8* resistance is largely overcome worldwide, we tailored the *B.g. tritici* diversity panel in order to increase the frequency of avirulent isolates. We did so by including a high proportion of isolates collected in the fertile crescent that were recently shown to exhibit the highest genetic diversity (Sotiropoulos *et al*., 2022). The GWAS diversity panel was subsequently phenotyped on three *Pm8* containing genotypes: the near-isogenic line ‘Kavkaz/4*Federation’, which is based on one of the earliest released commercial cultivars carrying the 1BL.1RS translocations ‘Kavkaz’, as well as two independent *Pm8* transgenic lines ‘Pm8#12’ and ‘Pm8#34’ described in (Hurni *et al*., 2013). 41 isolates, including the reference isolate ISR_7, showed an avirulent phenotype on the *Pm8* transgenic lines (i.e. no sporulation) and consistently exhibited reduced but residual sporulation on ‘Kavkaz/4*Federation’ indicating that endogenous expression levels of *Pm8* are not sufficient to provide complete resistance under laboratory conditions (Fig. 1a,b). In contrast, the remaining 38 isolates exhibited a virulent *Pm8* phenotype, efficiently infecting all three *Pm8*-containing genotypes (Fig. 1a,b). Genomic association between sequence polymorphisms (SNPs) and virulence/avirulence patterns on *Pm8* containing wheat lines identified two significantly associated SNPs, separated by 15’554 bp and mapping to an effector gene cluster on the short arm of *B.g. tritici* chromosome 11 (Fig. 1c-e). Best association was found for snp280882, located within the coding sequence of the *BgISR7-10067* effector gene. *BgISR7-10067* is part of effector gene family E003 (nomenclature of (Müller *et al*., 2019)) that also contains the recently identified *AvrPm17* (Fig. 1e, Fig. S1) (Müller *et al*., 2022). In a next step we compared the genomic region harboring *BgISR7-10067* in the *Pm8* avirulent isolate ISR_7 with the corresponding region in the genome assembly of isolate CHE_96224 (Müller *et al*., 2019), which exhibits a virulent phenotype on *Pm8* (Fig. 1a). Sequence comparison revealed high levels of co-linearity in this chromosomal region and only minor structural differences that do not affect any coding genes (Fig. 1e). Strikingly, the effector proteins encoded by *AvrPm8* candidate gene *BgISR7-10067* and *Bgt-50847*, its ortholog in isolate CHE_96224, differ by a single amino acid polymorphism (F43Y) which corresponds to the best associated SNP snp280882 in the GWAS analysis (Fig. 1e-f). Furthermore, BgISR7-10067 presents all the hallmarks of avirulence effectors in *B.g. tritici*, such as its small size (i.e. 107 amino acids), the presence of a signal peptide, a Y/FxC motif (Fig. 1f) and high expression levels during early infection stages (Fig. S2) (Bourras *et al*., 2015; Praz *et al*., 2017; Bourras *et al*., 2019; Hewitt *et al*., 2020; Müller *et al*., 2022). *BgISR7-10067* was therefore considered an excellent *AvrPm8* candidate gene.

**Figure 1:**
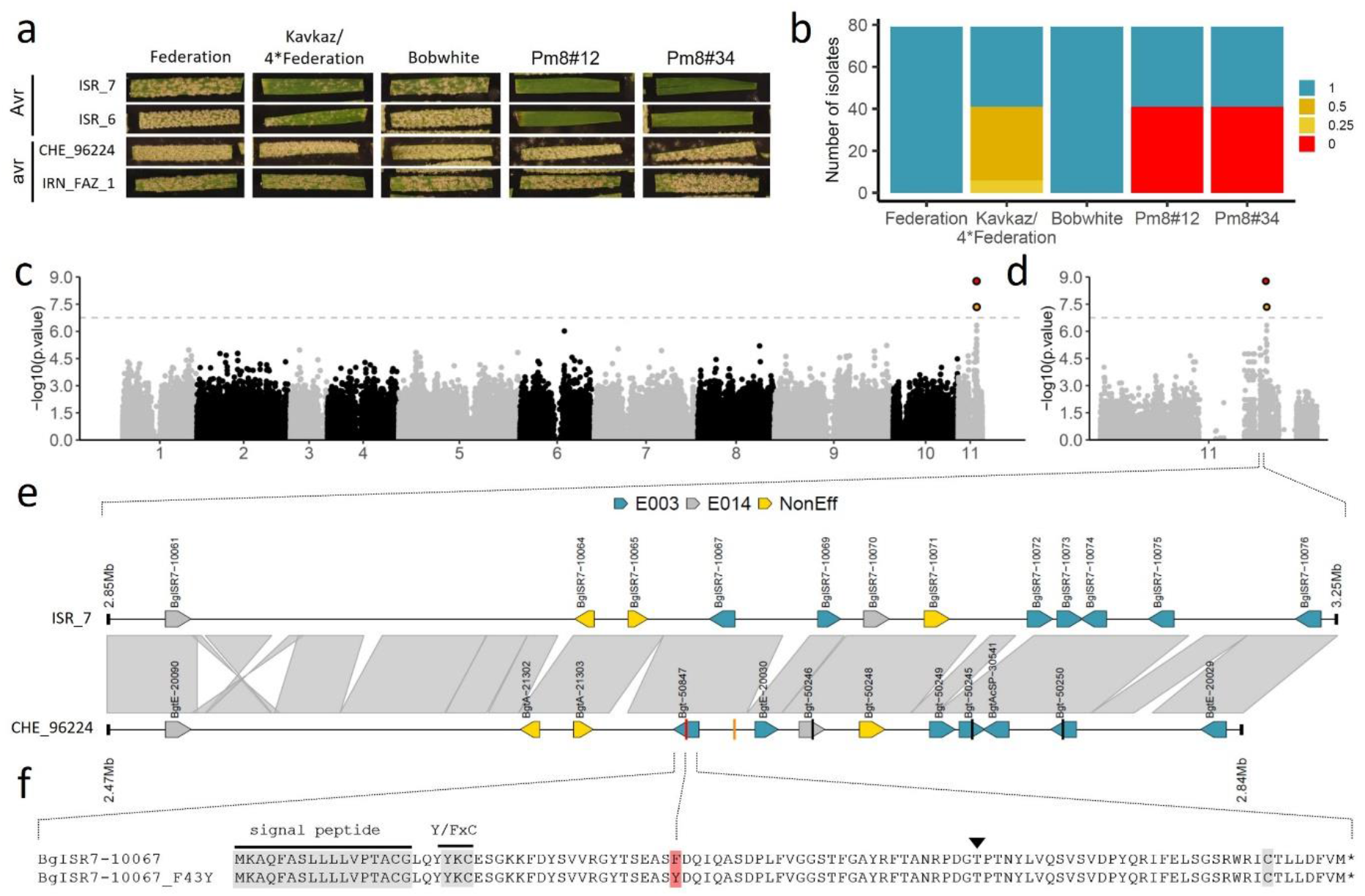
Identification of *AvrPm8* by GWAS. (**a**) Representative phenotypic spectrum of two avirulent (Avr) and two virulent (avr) *B.g. tritici* isolates on *Pm8* wheat lines ‘Kavkaz/4*Federation’ and two independent transgenic lines ‘Pm8#12’ and ‘Pm8#34’. Susceptible wheat cultivars ‘Federation’ and ‘Bobwhite’ served as controls. (**b**) Phenotypic distribution of 79 *B.g. tritici* isolates used for GWAS. Leaf coverage (i.e. virulence) was scored 8-10 days after infection using five categories ranging from 0 (0% leaf coverage) to 1 (100% leaf coverage). (**c,d**) Results of GWAS for avirulence on *Pm8*. Association between sequence polymorphisms and phenotype are depicted along the 11 chromosomes (**c**) or exclusively along chromosome 11 (**d**) of *Pm8* avirulent isolate ISR_7. Significance threshold at p<0.05 after Bonferroni correction is indicated as a dashed line. The two significantly associated SNPs on chromosome 11 are highlighted in red (best associated SNP) and yellow. (**e**) Sequence comparison of the genomic region harbouring the *AvrPm8* candidate gene *BgISR7-10067* in the genome assemblies of isolates ISR_7 (top track) and CHE_96224 (bottom track). Co-linearity is indicated between the two tracks (grey). Genes are indicated by arrows; gene models are not drawn to scale. Effector genes of effector family E003 (blue), effector family E014 (grey) are highlighted. Non-effector genes are depicted in yellow. The two significantly associated SNPs of the GWAS analysis are indicated by red (best associated SNP) and yellow lines. (**f**) Sequence alignment of the effector proteins encoded by *AvrPm8* candidate *BgISR7-10067* in ISR_7 and its orthologous gene *BgISR7-10067_F43Y* (*Bgt-50847*) in CHE_96224. Hallmarks of *B.g. tritici* effector proteins such as the predicted signal peptide the Y/FxC motif and a conserved cysteine at the C-terminus are indicated in grey. The best associated SNP of the GWAS analysis corresponding to the amino acid substitution F43Y is highlighted in red. The black arrowhead indicates the position of the intron.

In order to validate *AvrPm8*, we co-expressed *BgISR7-10067* together with *Pm8-HA* in *Nicotiana benthamiana* using transient *Agrobacterium* mediated overexpression (Bourras *et al*., 2019). To ensure efficient translation in planta, we codon-optimized all fungal effector constructs omitting the predicted signal peptide and fused them to a C-terminal FLAG epitope tag for protein detection by Western blotting. Co-expression of *BgISR7-10067* (hereafter referred to as *AvrPm8*) with *Pm8-HA* resulted in a strong cell-death response (hypersensitive response, HR) in *Nicotiana*, which was absent when either of the components was expressed alone (Fig. 2a,b), confirming *BgISR7-10067* as *AvrPm8*. In agreement with the virulent phenotype of isolate CHE_96224, its orthologous gene *Bgt-50847* (*AvrPm8_F43Y*) did not trigger any HR response upon co-expression with *Pm8* (Fig. 2a-c). Western blot analysis confirmed the efficient production of both FLAG-tagged AvrPm8 variants and Pm8-HA in *N. benthamiana*, indicating the F43Y mutation affects recognition of AvrPm8 by its cognate immune receptor Pm8 (Fig. 2d,e).

**Figure 2:**
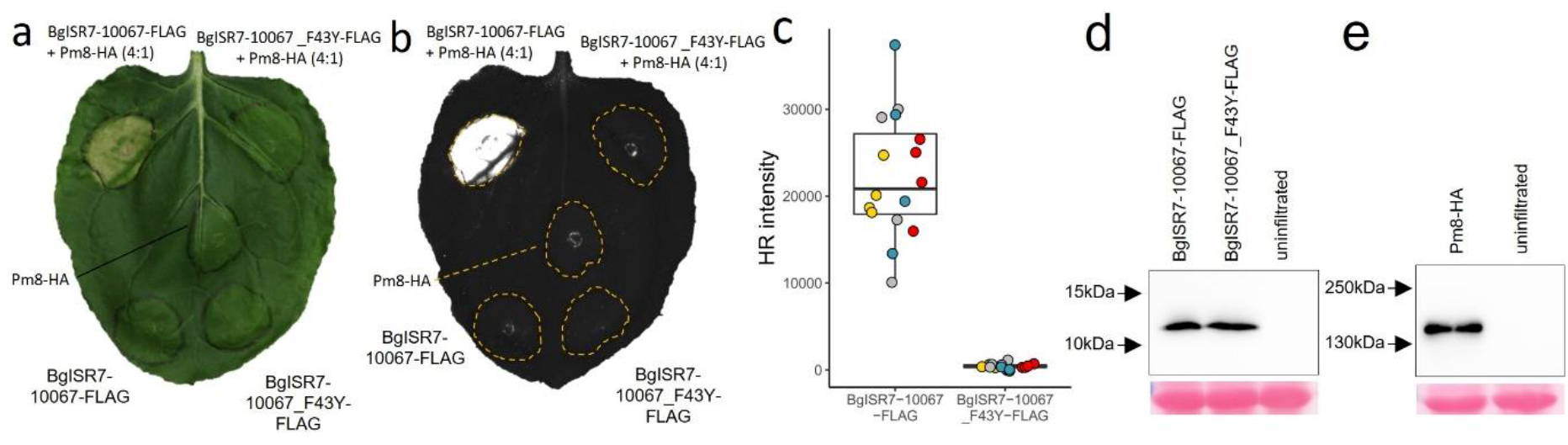
Functional validation of *AvrPm8* in *Nicotiana benthamiana*. (**a,b**) *Agrobacterium*-mediated transient expression in *Nicotiana benthamiana* of *BgISR7-10067-FLAG* (*AvrPm8*) or *BgISR7-10067_F43Y-FLAG* (*AvrPm8_F43Y*) co-infiltrated with (top) or without (bottom) *Pm8-HA*. Leaves were imaged at 5 days post inoculation using a normal camera (**a**) or using a Fusion FX imager system (**b**). Co-infiltrations were performed with a ratio of 4 (effector): 1 (Pm8-HA). Co-expression of *BgISR7-10067* with *Pm8* resulted in hypersensitive cell-death (HR) in four independent experiments with a total of n=16 leaves. Co-expression of *BgISR7-10067_F43Y* with *Pm8* or expression of any component alone did not result in HR (n=16). (**c**) Quantification of HR intensity in co-expression assay depicted in (**b**). Individual datapoints are color coded based on four independent experiments with n=4 leaves per experiment. (total n=16) (**d,e**) Detection of *N. benthamiana* expressed AvrPm8-FLAG and AvrPm8_F43Y-FLAG (**d**) or Pm8-HA (**e**) by anti-FLAG or anti-HA western blotting (top panel) or total protein Ponceau S staining (bottom panel). Black arrows indicate positions of protein size markers.

Given the rye origin of the *Pm8* resistance gene we searched for *AvrPm8* homologous genes in the rye infecting sublineage of *Blumeria*, *Blumeria graminis* f.sp. *secalis* (*B.g. secalis*). We hypothesized that based on its host range and divergence from *B.g. tritici* approximately 200’000 years ago (Menardo *et al*., 2016), *B.g. secalis* should have been exposed to the *Pm8* resistance specificity over a longer evolutionary time frame than *B.g. tritici*. In all tested *B.g. secalis* isolates (5) we found an *AvrPm8* homolog (*AvrPm8_Bgs*), encoding for a protein with 13 amino acid differences compared to AvrPm8 (Fig. 3a). Interestingly, the *AvrPm8* homologous gene in *B.g. secalis* harbored the identical sequence polymorphism found in *B.g. tritici*, leading to the amino acid substitution F43Y. Similar to AvrPm8_F43Y, co-expression of AvrPm8_Bgs with Pm8-HA in *N. benthamiana* did not result in an HR response (Fig. 3b). The efficient production of AvrPm8_Bgs protein in *N. benthamiana* (Fig. 3c), indicated that AvrPm8_Bgs indeed evades Pm8 recognition, at least in part due to its F43Y mutation. Consistent with our findings in *N. benthamiana*, *B.g. secalis* isolates exhibited full virulence on *Pm8* containing rye cultivars ‘Petkus’, the donor cultivar of the 1BL.1RS translocation (Lein, 1975), and ‘Lo7’ an inbred rye line which was recently shown to contain *Pm8* (Rabanus-Wallace *et al*., 2021) (Fig. 3d). These findings further corroborate the importance of the F43Y substitution in *AvrPm8* for the evasion of *Pm8* recognition and indicate the underlying DNA polymorphism to be relatively ancient.

**Figure 3:**
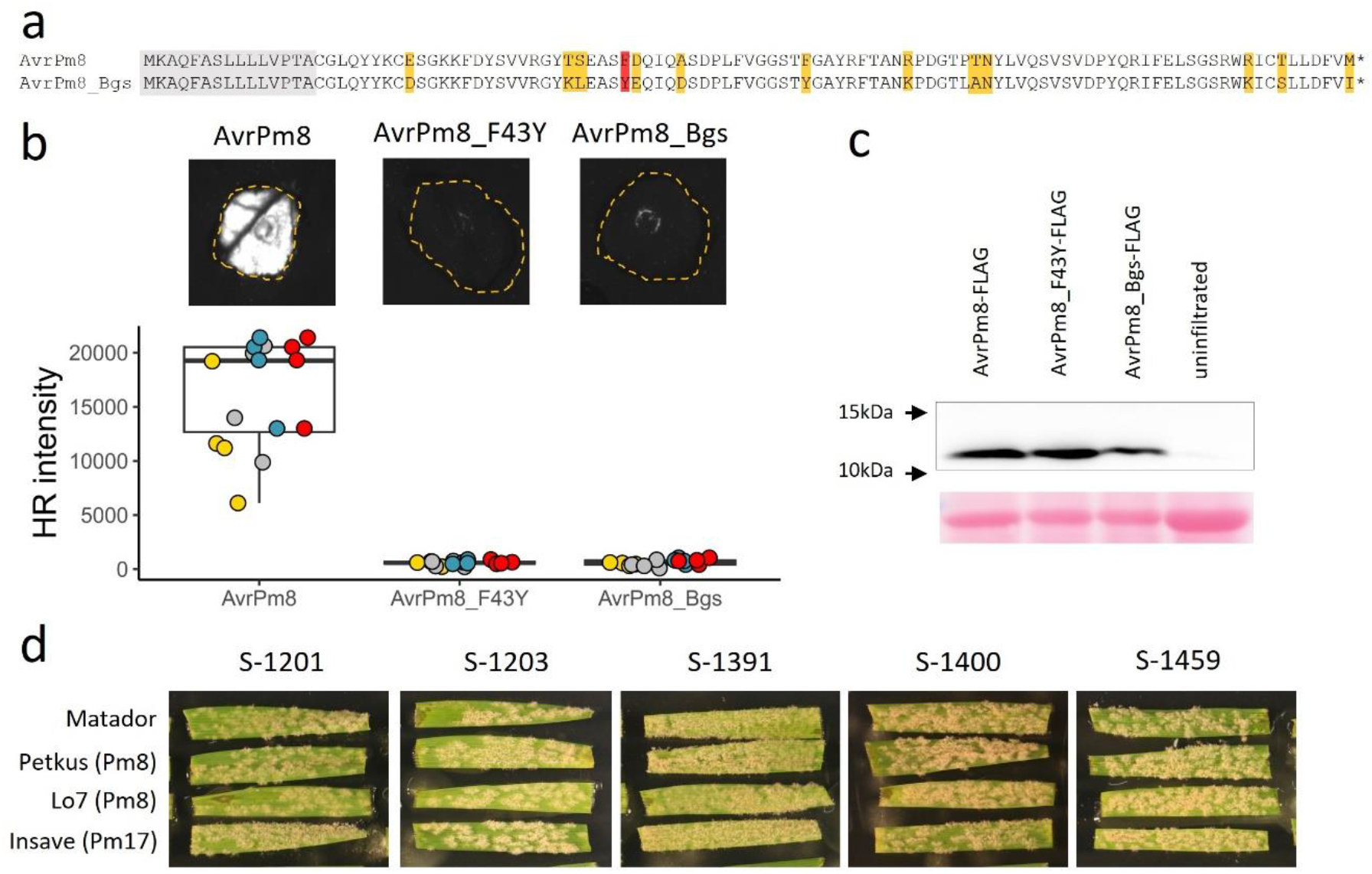
The *AvrPm8* homologous gene in *B.g. secalis* is not recognized by *Pm8*. **(a)** Protein sequence alignment of AvrPm8 and its *B.g. secalis* homolog AvrPm8_Bgs. Predicted signal peptide (gray), F43Y substitution (red) and additional polymorphic residues (yellow) are highlighted. **(b)** *Agrobacterium*-mediated transient co-expression of AvrPm8-FLAG, AvrPm8_F43Y-FLAG and AvrPm8_Bgs-FLAG with Pm8-HA. Leaves were imaged at 5 days post inoculation using a Fusion FX imager system (top panel). Co-infiltrations were performed with a ratio of 4 (effector) : 1 (Pm8-HA). Bottom panel: quantification of HR intensity in co-expression assay depicted in top panel. Individual datapoints are color coded based on four independent experiments with n=4 leaves per experiment (total n=16). **(c)** Detection *N.benthamiana* expressed AvrPm8-FLAG, AvrPm8_F43Y-FLAG and AvrPm8_Bgs-FLAG by anti-FLAG western blotting (top panel) or total protein Ponceau S staining as a loading control (bottom panel). Black arrows indicate positions of protein size markers. **(d)** Virulence phenotype of five *B.g. secalis* isolates on rye cultivars ‘Petkus’ and ‘Lo7’ both carrying a *Pm8* gene, rye cultivar ‘Insave’ carrying *Pm17* and ‘Matador’ as a susceptible control. Pictures were taken 10 days after infection.

In order to get a more in-depth view on the resistance breakdown of *Pm8* in wheat, we performed extensive haplotype mining of the *AvrPm8* gene in a global collection of 219 *B.g. tritici* isolates (Sotiropoulos *et al*., 2022). In addition to the above described *AvrPm8* and *AvrPm8_F43Y* variants, we identified a single synonymous mutation and 17 sequence polymorphisms that impact the open reading frame (ORF) of the *AvrPm8* gene in various ways (Fig. 4a). This included an additional point mutation affecting the crucial amino acid position F43 (F43L) and numerous mutations resulting in a premature stop codon (Q4STOP, K28STOP, D30STOP, S32STOP, L77STOP, R96STOP). Additionally, we identified three independent mutations disrupting the start codon ATG (ATG>GTG, ATG>AAG, ATG>ATC) (Fig. 4a). Furthermore, we identified four independent mutations affecting both terminal dinucleotides (i.e. splice acceptor and donor sites) of the single intron found in the *AvrPm8* gene (Fig. 4a). Mutations in the highly conserved terminal dinucleotides (GT-AG) have been found to disrupt mRNA maturation and result in exon skipping or retention of the intron (see Supplementary Note S1). We hypothesized that the disruption of splicing of the *AvrPm8* mRNA could represent a gain-of-virulence mechanism. Indeed, analysis of RNA sequencing data from GBR_JIW2, an isolate carrying a mutation in the 5’ dinucleotide leading to the splice sites GA-AG (instead of GT-AG), confirmed splicing of the intron to be largely abolished (Fig. 4a, Fig. S3, Supplementary Note S1). Ribosomal translation of unspliced *AvrPm8* mRNA would therefore lead to a premature stop codon upon translation of the intron and consequentially a truncated protein. In addition to all above-mentioned single nucleotide polymorphisms affecting the *AvrPm8* gene, we also found two independent, large deletion events of 14 and 43 kb, both encompassing the entire *AvrPm8* gene (Fig. 4a, Fig. S4).

**Figure 4:**
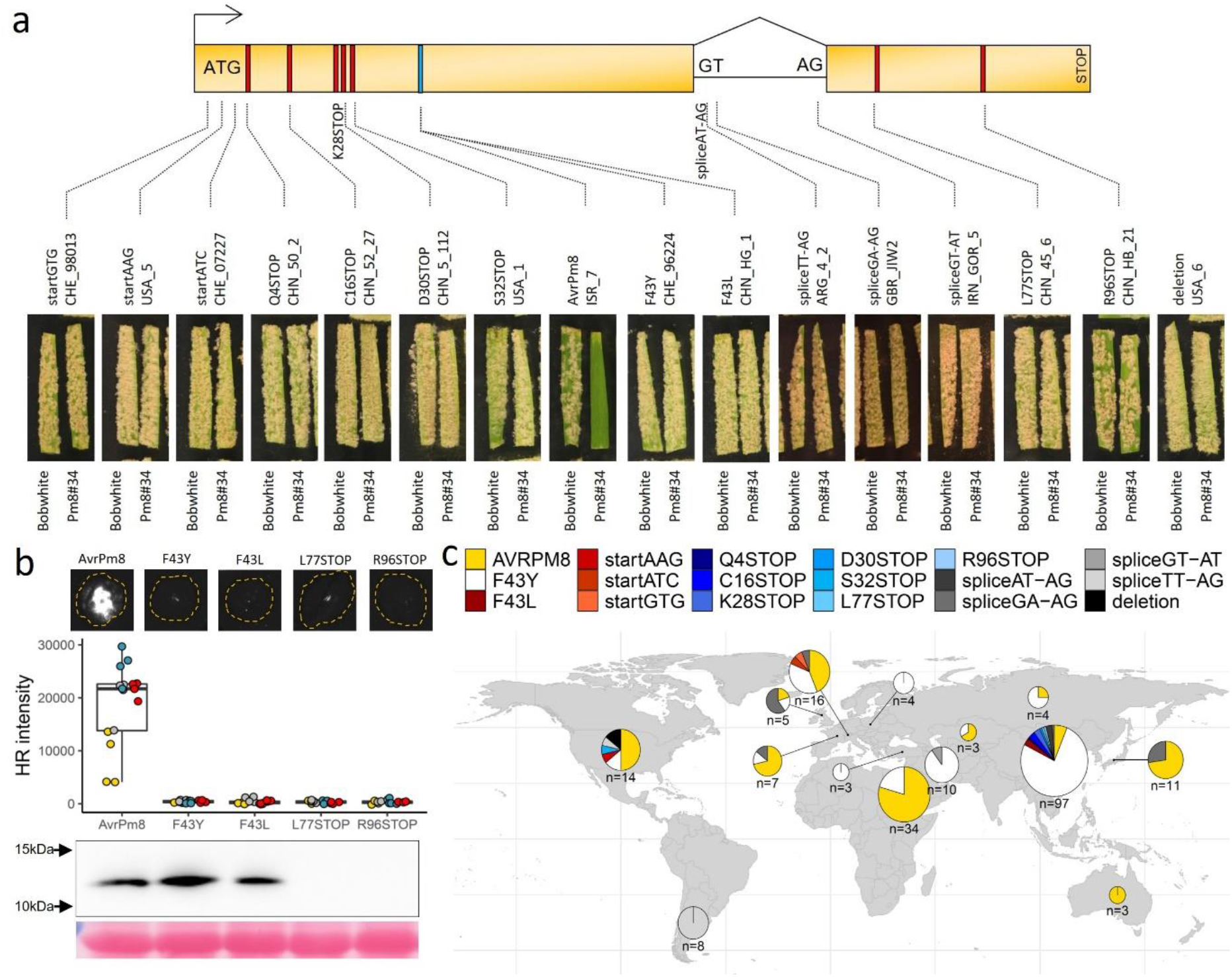
*AvrPm8* haplovariant mining in a global *B.g. tritici* population reveals 17 gain-of-virulence mutations. (**a**) Schematic gene model (toxp) and phenotypes (bottom) of isolates with the identified gain-of-virulence mutations in the *AvrPm8* effector gene. The two exons of *AvrPm8* are depicted as yellow boxes with gene orientation indicated by an arrow. Start codon, splice sites, and stop codon are indicated. Mutations leading to an amino acid change (blue), or a premature stop codon (red) are indicated by a coloured line. For each mutation the virulence phenotype of a representative isolate on the *Pm8* transgenic line Pm8#34 is depicted. Susceptible cultivar ‘Bobwhite’ serves as a control. The phenotype of the avirulent isolate ISR_7 is shown as comparison (middle). Additional phenotyping data for each isolate including phenotypes on the near-isogenic line ‘Kavkaz/4*Federation’ and an independent *Pm8* transgenic line Pm8#12 are depicted in Fig. S5. For two mutations (K28STOP and spliceAT-AG) we were unable to find a representative isolate in our living *B.g. tritici* collection. (**b**) *Agrobacterium* mediated co-expression of genes encoding AvrPm8-FLAG, AvrPm8_F43Y-FLAG, AvrPm8_F43L-FLAG, AvrPm8-R96STOP-FLAG or AvrPm8-L77STOP-FLAG together with Pm8-HA in *N. benthamiana*. Leaves were imaged at 5 days post inoculation using a Fusion FX imager system (top panel). Co-infiltrations were performed with a ratio of 4 (effector) : 1 (Pm8-HA). Middle panel: quantification of HR intensity in co-expression assay depicted in top panel. Individual datapoints are color coded based on four independent experiments with n=4 leaves per experiment (total n=16). Detection of *N. benthamiana* expressed AvrPm8 constructs by anti-FLAG western blotting (bottom panel) and total protein Ponceau S staining as a loading control. Black arrows indicate positions of protein size markers. (**c**) Worldwide distribution of identified *AvrPm8* haplovariants. Haplovariants are depicted in different colors according to the legend above the world map. Size of pie charts are scaled as log10(n) where n is the number of isolates per region.

In order to understand the impact of individual mutations found in *AvrPm8*, we tested representative isolates for each mutation on *Pm8* containing wheat lines ‘Kavkaz/4*Federation’ and the two transgenic *Pm8* lines, wherever isolates were available in our living *B.g. tritici* collection (15 out of 17 mutations). For all *AvrPm8* mutations tested, the affected isolates exhibited a fully virulent phenotype on *Pm8* wheat, indicating that all *AvrPm8* haplotypes apart from the ISR7 haplotype, represent virulence alleles (Fig. 4a, Fig S5). To verify this finding in *N. benthamiana*, we co-expressed *AvrPm8_F43L*, as well as the two longest variants with premature stop codons, *AvrPm8_L77STOP* and *AvrPm8_R96STOP*, together with *Pm8* in the *Nicotiana* system. Consistent with the phenotype of corresponding isolates on *Pm8* wheat, none of these AvrPm8 variants was recognized by Pm8 (Fig. 4b). Western blot analysis revealed efficient production of AvrPm8_F43L in *Nicotiana*, further pinpointing at the crucial role of the phenylalanine (F) at position 43 for Pm8 recognition (Fig. 4b). In contrast, both truncated AvrPm8 variants (L77STOP and R96STOP) were undetectable in protein extracts from Nicotiana, indicating the truncations result in an unstable AvrPm8 protein (Fig 4b). Such instability also has to be assumed for all other premature stop codon inducing mutations (Q4STOP, K28STOP, D30STOP, S32STOP), resulting in an even shorter open reading frame.

We then analyzed the worldwide distribution of the avirulent *AvrPm8* variant and any of the above-described gain-of-virulence mutations. Several patterns became apparent. Firstly, the recognized *AvrPm8* variant is found in most subpopulations worldwide (Fig. 4c). Given the fact *Pm8* resistance is largely broken *AvrPm8* was surprisingly frequent in populations in Central Europe, the US and particularly in Israel, Japan, and Australia (Fig. 4c). While some of these regions suffer from undersampling and interpretations should be drawn with caution, the dominance of *AvrPm8* in the well-covered Israeli subpopulation is striking. We hypothesize that the Israeli population was exposed to a less severe *Pm8* dependent selection pressure due to the existence of *Pm8*-free wild wheat progenitors growing naturally in this region. Similar effects might be in place in other regions of the world where *Pm8* containing wheat cultivars did not reach a high popularity. The only virulent variant with worldwide distribution is the *AvrPm8_F43Y* allele. It represents the most common *AvrPm8* haplotype found worldwide and dominates in frequency in many subpopulations on the Eurasian continent and particularly in China (Fig. 4c). Based on its widespread occurrence and the fact that an identical mutation is found also in *B.g. secalis* (Fig. 3a) we hypothesize that the *AvrPm8_F43Y* virulence allele is ancient and precedes the introgression of *Pm8* into wheat. It has probably increased in frequency within the *B.g. tritici* populations of regions with prevalent *Pm8* use such as China or Eastern Europe (Fig. 4c). In contrast to the broadly found *AvrPm8* and *AvrPm8_F43Y* variants, all other *AvrPm8* haplovariants were found only in few isolates often with geographically very restricted occurrence (Fig. 4c). For example, the AvrPm8_F43L haplovariant was found exclusively in China, the three identified start codon mutations were found in single isolates either in Switzerland (startGTG, startATC) or the US (startAAG) and the multiple variants with premature stop codons were unique to China (Q4STOP, K28STOP, D30STOP, L77STOP, R96STOP) or the US (S32STOP). In particular the US and Chinese *B.g. tritici* populations harbored many unique gain-of-virulence mutations of *AvrPm8*. Our findings indicate that, apart from the F43Y substitution, gain-of-virulence mutations represent relatively recent events, likely as a consequence of local selection pressure exerted by the high prevalence of *Pm8* wheat.

## Discussion

Genomic analyses of *B.g. tritici* and *Blumeria graminis* f. sp. *hordei* (barley powdery mildew) revealed that a high proportion of effector genes encode for small, secreted proteins with a predicted RNAse-like structure (Pedersen *et al*., 2012; Menardo *et al*., 2017; Seong & Krasileva, 2022). Several RNAse-like effectors have furthermore been implicated in virulence processes of *Blumeria* (Zhang *et al*., 2012; Pliego *et al*., 2013; Ahmed *et al*., 2015; Ahmed *et al*., 2016; Pennington *et al*., 2019). It is therefore not surprising that up to date all identified avirulence proteins in *B.g. tritici* and *B.g. hordei* fall into this effector category. The newly identified AvrPm8 is no exception to this rule and exhibits common features like small size, presence of a signal peptide, a Y/FxC motif, and a conserved cysteine towards the C-terminus. Like AvrPm17, it furthermore belongs to effector family E003, previously shown to contain effectors with a predicted RNAse-like structure (Saur *et al*., 2019; Bauer *et al*., 2021; Müller *et al*., 2022). The recently identified *AvrPm3^a2/f2^*, *AvrPm3^b2/c2^*, *AvrPm3^d3^* and *AvrPm17*, are recognized by NLRs encoded by the *Pm3* allelic series and the Pm3 homologous rye NLR Pm17, respectively. They all exhibited exceptionally high expression levels during early stages of the infection process, indicating an important function in the establishment of a successful infection (Bourras *et al*., 2015; Bourras *et al*., 2019; Müller *et al*., 2022). Again, *AvrPm8* shows a similar trend by ranking among the 5% highest expressed genes during infection (Fig. S2).

Haplovariant mining approaches for *AvrPm3* and *AvrPm17* combined with functional characterization of the diverse natural variants have led to a detailed understanding of gain-of-virulence mechanisms and associated resistance breakdown of the *Pm3* allelic series and its rye homolog *Pm17*. Interestingly, *AvrPm3* and *AvrPm17* gain-of-virulence mutations were found to rely on single amino acid polymorphisms and copy number variation in some cases, while preserving at least one functional ORF of the effector gene in each case (McNally *et al*., 2018; Bourras *et al*., 2019; Müller *et al*., 2022). These findings contrast with gain-of-virulence mutations in recently identified avirulence genes *AvrSr27*, *AvrSr35* and *AvrSr50* of the biotrophic wheat stem rust pathogen, which frequently involved avirulence gene deletion, transposable element insertion or drastic expression polymorphisms (Chen *et al*., 2017; Salcedo *et al*., 2017; Upadhyaya *et al*., 2021). The conservation of an intact ORF in virulence alleles of *AvrPm17* and *AvrPm3* genes, was interpreted as indications for the counterselection of deleterious mutations due to the importance of the effector function for *B.g. tritici* virulence. Haplovariant mining in a worldwide *B.g. tritici* diversity panel for *AvrPm8* also identified two single amino acid polymorphisms leading to gain-of-virulence. Strikingly, the F43Y and F43L mutations affected the same amino acid, indicating an important function of phenylalanine at position 43 for recognition by Pm8 (Fig. 4a, b). Whether mutations in other regions of the AvrPm8 protein would not lead to gain-of-virulence or would be disadvantageous to the pathogen remains to be determined. The presence of 12 additional amino acid polymorphisms in the AvrPm8 homolog found in *B.g. secalis* (Fig. 3a) indicates that polymorphisms throughout the AvrPm8 protein can be tolerated. It is possible that missense mutations in other regions of the AvrPm8 protein would result in only partial gain-of-virulence phenotypes on *Pm8* which, due to the broad application of *Pm8*, would be quickly outcompeted by fully virulent *AvrPm8* haplovariants. Indeed, partial virulence phenomena have been observed for several naturally occurring amino acid polymorphisms in the closely related *AvrPm17* (Müller *et al*., 2022). We hypothesize that expanding haplovariant mining efforts would reveal additional amino acid polymorphisms, even though they might be less frequent as compared to the *AvrPm3*’s and *AvrPm17*.

In contrast to the conservation of intact open reading frames in *AvrPm3* and *AvrPm17* virulence alleles, we found strong convergent evolutionary trends for the occurrence of more drastic virulence mutations to the *AvrPm8* gene. Not only did we find evidence for numerous independent mutation events affecting the same structural components of the *AvrPm8* gene such as the start codon or splice sites, but also most of the identified gain-of-virulence mutations lead to significant truncations or the complete destruction of the *AvrPm8* ORF. Strikingly, the natural diversity of *AvrPm8* virulence alleles found in the worldwide *B.g. tritici* population strongly resembled the outcome of an artificial EMS mutagenesis screen for gain-of-virulence in *AvrSr35*, which identified 15 virulence inducing mutations involving premature stop codons (12), splice site mutations (1) and two independent mutations affecting the same amino acid (Salcedo *et al*., 2017). Even though the virulence function of the AvrPm8 effector is currently unknown, given the frequency of deleterious mutations to the AvrPm8 ORF under natural conditions, we hypothesize that AvrPm8 effector activity, in contrast to AvrPm17 and AvrPm3 effectors, is dispensable without major fitness costs in *B.g. tritici*. We therefore argue that the diverse deterioration of the *AvrPm8* gene observed in *B.g. tritici* is a consequence of the dispensability of AvrPm8 effector function in combination with the strong selection pressure exerted by the broad and frequent use of the *Pm8* resistance gene as part of the 1BL.1RS translocation in wheat.

The deployment of 1BL.1RS wheat cultivars in larger agricultural settings is first reported in 1960’s followed by its integration into national and international breeding programs and associated increase in planted acreage in the following decades (Lukaszewski, 1990; Rabinovich, 1998; Graybosch, 2001; Zhou *et al*., 2007). In many countries the *Pm8* resistance breakdown was however reported within few years of 1BL.1RS deployment (Bennett, 1984; Streckeisen, 1985; Namuco *et al*., 1987; Heun & Friebe, 1990; Purnhauser *et al*., 2011). Given the broad application and reported near-complete *Pm8* breakdown, we were surprised by the prevalence of the avirulent *AvrPm8* haplovariant in specific local *B.g. tritici* populations such as Australia, Israel, Japan or Central Europe. We hypothesize that this reflects consequences of local breeding decisions (i.e. low frequency of *Pm8* cultivars) or, in the case of Israel, the availability of sympatrically growing, *Pm8*-free, wild wheat relatives such as wild emmer known to allow growth of *B.g. tritici* (Ben-David *et al*., 2016).

Interestingly we found only a single virulence allele, *AvrPm8_F43Y* with worldwide occurrence. Its widespread distribution and the presence of an identical nucleotide polymorphism in the related *B.g. secalis* sublineage, suggests the F43Y mutation is ancient and likely precedes *Pm8* introgression from rye to wheat. This is reminiscent of the ancient genetic variation identified in *AvrPm17*, including multiple gain-of-virulence mutations, explaining the quick breakdown of the *Pm17* resistance gene introgressed into wheat (Müller *et al*., 2022). Interestingly *AvrPm8_F43Y* dominated *B.g. tritici* population in many regions where the 1BL.1RS translocation dominantly impacted local breeding programs, and the *Pm8* resistance was reported to be completely broken, such as Eastern Europe and in particular China (Rabinovich, 1998; Zhou *et al*., 2007; Purnhauser *et al*., 2011; Zeng *et al*., 2014). Especially the Chinese *B.g. tritici* subpopulation analyzed in this study exhibited striking patterns of a strong selection for virulence on *Pm8* lines consistent with previous reports (Zhou *et al*., 2007; Zeng *et al*., 2014). Among the 97 analyzed isolates only six carried the avirulent *AvrPm8* haplovariant, with the remaining isolates carrying either *AvrPm8_F43Y* (74) or one of eight locally occurring gain-of-virulence variants. The near complete absence of the avirulent *AvrPm8* is reminiscent of a recent study in a global population of Septoria leaf blotch which found absence of avirulent forms of *AvrStb6* in modern Septoria isolates due to widespread use of *Stb6* wheat in recent years (Stephens *et al*., 2021). The Chinese *B.g. tritici* population furthermore exemplifies the observed global trend for rare, locally restricted gain-of-virulence innovations that are represented by mostly deleterious mutations to the *AvrPm8* open reading frame. Based on their restricted distribution we hypothesize that these *AvrPm8* virulence variants represent recent mutational events that occurred and propagated locally upon exposure of a *B.g. tritici* subpopulation to *Pm8*.

It has been assumed, that the near simultaneous breakdown of powdery mildew resistance genes such as *Pm17* or *Pm8* in wheat growing areas worldwide can, at least partially, be explained by the ability of powdery mildew ascospores to travel large distances within only few growing seasons thereby allowing the efficient spread of new gain-of-virulence alleles (Limpert *et al*., 1999; McDonald & Linde, 2002). However, a recent population genomics analysis by Sotiropoulos et al. (Sotiropoulos *et al*., 2022), making use of the same *B.g. tritici* diversity panel used for this study, found strong associations between genetic proximity and geographic origin, and convincingly attributed the exchange of mildew genetic material over long distances (i.e. continents) to human activities in historic times followed by hybridization events. These findings are consistent with our hypothesis of frequent, locally restricted *de novo* gain-of-virulence mutations in *AvrPm8* leading to the seemingly simultaneous breakdown of *Pm8* resistance worldwide.

The quick breakdown of *Pm8* by convergent evolution in powdery mildew contrasts the breakdown of the more durable *Sr31* resistance gene residing on the same 1BL.1RS rye translocation. Even though the underlying resistance mechanism of *Sr31* is not known, the emergence of the *Sr31* virulent race Ug99 in Uganda could be attributed to a single evolutionary event involving nuclear exchange during somatic hybridization (Li *et al*., 2019) and subsequent stepwise spread of the Ug99 lineage to countries in Africa and the Middle East (Singh *et al*., 2011). The differences in the breakdown of *Pm8* and *Sr31* exemplify the different evolutionary dynamics involved in resistance gene breakdown, likely influenced not only by the molecular mode of action of the resistance gene but also by the lifestyles, genetic diversity, and genome stability of the involved fungal pathogens. We therefore consider it crucial to expand the study of fungal avirulence gene dynamics in various wheat pathogens in order to understand emerging patterns of resistance gene breakdown and consequentially allow informed breeding decisions towards more durable resistance.

## Methods

### Plant material, fungal isolates, and virulence phenotyping

The ‘Kavkaz/4*Federation’ *Pm8* near-isogenic line has been previously described in (Hurni *et al*., 2013). The two independent *Pm8* transgenic lines carrying the complete genomic sequence of the *Pm8* resistance gene under a maize ubiquitin promoter have been selected as best performing lines in the study of (Hurni *et al*., 2013). The *Pm8* rye cultivar ‘Petkus’, serving as the original donor for 1BL.1RS translocation in wheat, and the inbred rye line ‘Lo7’ have been previously shown to contain the *Pm8* resistance gene (Hurni *et al*., 2013; Rabanus-Wallace *et al*., 2021). The rye cultivar ‘Insave’ carrying the *Pm17* resistance gene has been described in (Singh *et al*., 2018).

Information about the collection of 219 *B.g. tritici* and 5 *B.g. secalis* isolates is summarized in Supplementary Dataset 1 and, with the exception of 10 newly sequenced Iranian isolates as part of this study, is described in great detail in (Sotiropoulos *et al*., 2022). *B.g. tritici* and *B.g. secalis* isolates were maintained clonally on leaf segments of susceptible bread wheat cultivar ‘Kanzler’ and susceptible rye cultivar ‘Matador’, respectively. For fungal propagation detached leaves were placed on 0.5% food grade agar (PanReac AppliChem) plates containing 4.23mM benzimidazole (Parlange *et al*., 2011).

For *B.g. tritici* isolates *Pm8* virulence phenotypes were assessed on the near-isogenic line ‘Kavkaz/4*Federation’ and two independent *Pm8* transgenic lines Pm8#12 and Pm8#34 (Hurni *et al*., 2013) with the susceptible control ‘Federation’ and ‘Bobwhite’, respectively. For *B.g. secalis* isolates *Pm8* virulence phenotypes were assessed on rye cultivars ‘Petkus’ and ‘Lo7’ (both with *Pm8*), ‘Insave’ (*Pm17*) and on the susceptible control ‘Matador’. Virulence scoring at 8-10 days after infection was performed on at least three biological replicates and based on qualitative assessment of mildew leaf coverage (MLC) in five categories: virulent (V): 100% MLC; intermediate/virulent (I/V): 75% MLC; intermediate (I): 50% MLC; avirulent/intermediate (A/I): 25% MLC; avirulent (A): 0% MLC. Phenotyping results are summarized in Supplementary Dataset 1.

### Bioinformatic datasets

The chromosome-scale assembly of isolate CHE_96224 (Bgt_genome_v3_16) is described in (Müller *et al*., 2019). The genome assembly of ISR_7 is described in (Müller *et al*., 2022). To resolve the chromosomes of the ISR_7 genome, the contigs of the ISR_7 assembly (available at the European nucleotide archive under accession number CAKMHR020000001 - CAKMHR020000032) were aligned against Bgt_genome_v3_16 using blastn (Camacho *et al*., 2009) and ordered according to the best blasthit (first criterium: evalue, second criterium: bitscore). The ordered assembly was subsequently polished using Illumina reads of ISR_7 previously published (SRA: SRX1140177, (Menardo *et al*., 2016). First, Illumina reads were quality trimmed using sickle (v1.33, https://github.com/najoshi/sickle) with options: pe -q 33 -l 40 and then aligned to the ISR_7 assembly using bowtie2 with the following parameters: −score-min L,−0.6,−0.25 (v2.3.4.1, (Langmead & Salzberg, 2012). Mappings files were processed with SAMtools (v1.7, (Li *et al*., 2009) sort, view, and rmdup commands followed by Picards (https://broadinstitute.github.io/picard/)) AddOrReplaceReadGroups command. Assembly polishing was done with pilon (v1.23, (Walker *et al*., 2014) with the -fix bases option. The final chromosome assembly is available at ENA accession number: CAKMHR020000000.1).

To create a draft annotation of the ISR_7 assembly, maker (v2.31.10, (Cantarel *et al*., 2008) was used with prot2genome option based on proteins of CHE_96224 (version Bgt_CDS_v4_23, available: https://zenodo.org/record/7018501) with repeat masking of the maker internal TE proteins, as well as the PTREP_2019 and nrTREP_2019 databases (available: https://trep-db.uzh.ch/). Subsequently, above-described steps were repeated with protein sequences from *B. g. hordei* isolates DH14 and RACE1 (GCA_900239735.1, GCA_900237765.1). Gene models from the second round were only added if the identified loci did not contain a gene model from the first round of annotation. The resulting draft annotation file is available at https://zenodo.org/record/6998719.

Whole-genome resequencing datasets used in this study have been described in (Sotiropoulos *et al*., 2022) unless indicated otherwise. Isolation, DNA extraction and sequencing of the 10 Iranian *B.g. tritici* strains newly described in this study was achieved as described by (Sotiropoulos *et al*., 2022). All whole-genome resequencing datasets are available from the SRA under project numbers indicated in Supplementary Dataset 1.

Raw RNA-sequencing reads of isolate CHE_96224 and GBR_JIW2 infecting wheat cultivar “Chinese Spring” at two days post infection (2dpi) are available at sequence read archive (SRA) under accession number SRP127348 (Praz *et al*., 2018). RNA-sequencing reads of isolate ISR_7 infecting wheat cultivar “Chinese Spring” at 2dpi were produced as described in (Praz *et al*., 2018) and sequenced on a NovaSeq, PE150 with 12Gb/20M reads total data output for each of three biological replicates (available from SRA under accession number PRJNA870298).

### Expression analysis

Expression analysis of ISR_7 was performed using salmon (v0.7.2 (Patro *et al*., 2017). First, CDS file of ISR_7 (available at https://zenodo.org/record/6998719) was index using the salmon index command. Subsequently, read counts per gene were calculated with the salmon quant command. Then count data was normalized using the calcNormFactors(method="TMM”) command from edgeR package (v3.38.4, (Robinson *et al*., 2010) and rpkm values per gene were calculated using the rpkm() command. Gene expression of all expressed genes (average rpkm value of three replicates >0) was plotted with a custom R script available at https://github.com/MarionCMueller/AvrPm8).

### Genome-wide association studies (GWAS)

For GWAS analysis, Illumina sequences of the 79 isolates (as indicated in Supplemental Dataset 1) were mapped against the ISR_7 genome assembly. First, sequences were quality trimmed using Trimmomatic (v0.38, (Bolger *et al*., 2014) using option LEADING:3 TRAILING 3 SLIDING WINDOW:4:20 MINLEN:50. Next, paired reads were aligned to ISR_7 assembly using bowtie2 (v2.2.9, (Langmead & Salzberg, 2012) with parameters −score-min L,−0.6,−0.25. Files were subsequently processed with SAMtools view, sort and rmdup commands (v1.6, (Li *et al*., 2009). Finally, files were processed using Picard (https://broadinstitute.github.io/picard/) with the ADDOrREplaceREADGroups command. SNPcall was performed with freebayes (v1.1.0-54-g49413aa, https://github.com/freebayes/freebayes) with -genotype quality and -p 1 options. The resulting vcf was filtered using vcftools (v0.1.5, (Danecek *et al*., 2011)) with the following parameters: --maf 0.05, --max-alleles 2 -min-alleles 2 -minDP10 -minGQ 40 -minQ 40 -max-missing 1 -remove-indels. The filtered vcf file was transformed to hapmap format using a custom perl script. GWAS was performed using GAPIT3 ((Wang & Zhang, 2021) using the following options: PCA.total =3, Model.selection = TRUE, model="GLM",kinship.algorithm = “VanRanden”. Scripts and dataset used to run the GWAS analysis are available at (https://github.com/MarionCMueller/AvrPm8). Manhattan plots were visualized with a custom R script available at https://github.com/MarionCMueller/AvrPm8).

### *AvrPm8* haplotype mining

For *AvrPm8* haplotype mining, whole-genome resequencing datasets were mapped to the *B.g. tritici* reference assembly of CHE_96224 (Bgt_genome_v3_16, (Müller *et al*., 2019)) using bwa (0.7.17-r1188, (Li & Durbin, 2009) as described in (Sotiropoulos *et al*., 2022) and visualized using the integrative genomics viewer IGV v2.8.6 (Robinson *et al*., 2011). *AvrPm8* haplotypes were defined by manual inspection for each isolate.

### Analysis of *AvrPm8* splicing

To analyze splicing of the *AvrPm8* mRNA, RNA-sequencing reads of isolates ISR_7 and GBR_JIW2 were aligned to the genome assembly of ISR_7 using STAR (v2.5.3, (Dobin *et al*., 2013) as follows: first, an index of the genome was created using the --genomeGenerate command. Secondly, RNAseq reads were aligned using with options -alignIntronMax 500, --outFilterMultimapNmax 5, --outFilterMismachNoverLmax 0.04 --outSAMtype BAM SortedByCoordinate. The resulting .bam files of three biological replicates were merged using the BamTools merge command (v2.5.1, https://github.com/pezmaster31/bamtools). Merged .bam-files were indexed using the SAMtools index command (v1.6, r(Li *et al*., 2009)) and visualized using the integrative genomics viewer (IGV) (v2.8.10, (Robinson *et al*., 2011)).

### Comparative genomics analysis of the *AvrPm8* locus

To analyze co-linearity of the *AvrPm8* locus between the two genome assemblies of ISR_7 and CHE_96224, the Chr-11 of both genome assemblies was compared using the nucmer command of the MUMmer program (v4.0.0rc1, (Marcais *et al*., 2018)). Subsequently the generated delta file was processed with the dnadiff command of the MUMmer suite. The resulting file out.1coords was visualized in R studio using the gggenomes package (https://github.com/thackl/gggenomes). Gene models in the locus were quality controlled manually, gene models that were erroneous or were not supported by RNA-seq reads were not depicted. Scripts and input files used to generate Fig. 1e are available at Github (https://github.com/MarionCMueller/AvrPm8)

### Phylogenetic tree

To define E003 effector family members in ISR_7, protein sequences of E003 family members of *B.g. tritici* reference isolates Bgt_96224 (Müller *et al*., 2022) were blasted against the protein sequences of ISR_7 (available at https://zenodo.org/record/6998719) using the blastp command from BLAST+ v2.6.0+, (Camacho *et al*., 2009) with e-value cutoff of 10e-50. Protein sequences were aligned using muscle algorithm implemented in MEGA-X software (v10.0.5, (Kumar *et al*., 2018)). The sequence of BgtE-20002 (AVRPMB2/C2) was used as outgroup. Phylogenetic tree was constructed using FastTree software (v2.1.11,(Price *et al*., 2010)) with default parameters. FigTree (v1.4.4, https://github.com/rambaut/figtree/releases/tag/v1.4.4) was used to visualize the tree. Protein sequences of E003 family members and the raw tree file is available at (https://github.com/MarionCMueller/AvrPm8).

### Verification of *AvrPm8* deletion by PCR

The extent of the *AvrPm8* gene deletion in USA_6 and USA_Ken_4_3 was estimated based on whole-genome resequencing data (see *AvrPm8* haplotype mining). For each deletion, a primer pair flanking the possibly deleted region was designed (for primer sequences see Supplementary Table S1). Genomic DNA was extracted from fungal spores using the chloroform and CTAB based extraction procedure described by (Bourras *et al*., 2015). PCR amplification spanning the gene deletion was achieved using Phusion High-Fidelity DNA Polymerase (New England Biolabs) according to the manufacturers protocol and visualized on 1% agarose gel supplied with ethidium bromide.

### Cloning of expression constructs

A gateway system compatible entry clone carrying the entire *Pm8* genomic sequence fused to a C-terminal Myc-tag (pENTR-Pm8-myc) has been described by (Hurni *et al*., 2014). For this study, the myc-epitope tag was replaced by a hemagglutinin (HA) epitope (creating pENTR-Pm8-HA) by PCR based site-directed mutagenesis (SDM) using non-overlapping primers listed in Supplementary Table S1 and Phusion High-Fidelity DNA Polymerase (New England Biolabs). Subsequently, the linear PCR product was phosphorylated using T4 polynucleotide kinase (New England Biolabs) and ligated with T4 DNA Ligase (New England Biolabs) according to the manufacturer.

The coding sequence, omitting the signal peptide as predicted by SignalP4.0 (Petersen *et al*., 2011), of all *AvrPm8* effector variants was codon-optimized for *N. benthamiana* using the tool provided by Integrated DNA technologies (https://eu.idtdna.com), C-terminally fused to a FLAG epitope tag, and synthesized with gateway compatible attL sites by our commercial partner BioCat GmbH (https://www.biocat.com). Sequence information of all constructs produced by gene synthesis can be found in Supplementary Dataset 2.

All gateway compatible entry clones described above were mobilized into the binary expression vector pIPKb004 (Himmelbach *et al*., 2007) using Gateway LR clonase II (Invitrogen) according to the manufacturer and subsequently transformed into *Agrobacterium tumefaciens* strain GV3101 using a freeze-thaw transformation protocol (Weigel & Glazebrook, 2006).

### Avr-R co-expression and HR quantification in *N. benthamiana*

*Agrobacterium tumefaciens* mediated expression of resistance and effector genes was achieved following the detailed protocol of (Bourras *et al*., 2019). For co-expression, *Agrobacteria* carrying the effector or resistance gene were mixed in a ratio of 4:1 directly prior to infiltration. HR development was assessed 4-5 days after infiltration using a Fusion FX imaging System (Vilber Lourmat, Eberhardzell, Germany) and quantified using Fiji (Schindelin *et al*., 2012) as described by (Bourras *et al*., 2019).

### Western blotting

To test for protein production in *N. benthamiana*, single constructs were expressed as described above and infiltrated plant tissue harvested 2 days after *Agrobacterium* infiltration. Protein extraction was achieved as described in (Bourras *et al*., 2019). Protein extracts were separated on homemade 4-20% gradient SDS polyacrylamide gels and blotted to a nitrocellulose membrane (Amersham Protran 0.2 μm NC) using a Trans-Blot SD Semi-Dry Transfer Cell (BioRad). Blotting efficiency was assessed by staining total protein with Ponceau S. For the detection of Pm8-HA anti-HA-HRP antibody (rat monoclonal, clone 3F10, Roche) was used at a dilution of 1:3000. For detection of FLAG tagged AvrPm8 variants, anti-FLAG-M2-HRP (mouse monoclonal, clone M2, Sigma-Aldrich) was used at a dilution of 1:3000. Peroxidase chemiluminescence was detected using a Fusion FX imaging System (Vilber Lourmat, Eberhardzell, Germany) and SuperSignal West Femto HRP substrate (Thermo Scientific) for FLAG tagged effectors or WesternBright ECL HRP substrate (Advansta) for Pm8-HA.

## Data availability

Illumina sequences used in this study are available at the short read archive under accession number PRJNA290428 and PRJNA625429. RNA-sequencing reads are available under accession number PRJNA427159 and PRJNA870298. Genome assembly of CHE_96224 and ISR_7 available under European nucleotide archive (ENA) accession number PRJEB28180 and PRJEB41382, respectively. Draft annotation of ISR_7 is available at https://zenodo.org/record/6998719.

## Acknowledgements

This study was supported by Swiss National Science Foundation grants 310030B_182833 and 310030_204165 to BK and the University Research Priority Program (URPP) ‘Evolution in Action’ of the University of Zurich. We would like to thank Andres Gordillo from KWS for providing the ‘Lo7’ seeds as well as Helen Zbinden and Esther Jung for the maintenance of powdery mildew isolates.

## Author contributions

L.K., M.C.M and B.K. designed the research. L.K., M.C.M and B.K. wrote the manuscript. L.K. performed the experiments. M.C.M, L.K., A.G.S and J.G. performed bioinformatic analyses. L.K. and M.C.M analysed data. M.R. collected mildew isolates.

## Supporting information

**Figure S1: Phylogenetic analysis of effector family E003 in *B.g. tritici* isolate ISR_7.**

**Figure S2: *AvrPm8* (*BgISR7-10067*) is among the top 5% of expressed genes in *B.g. tritici* isolate ISR_7.**

**Figure S3: Splice site mutations in isolate GBR_JIW2 abolish splicing of *AvrPm8* (*BgISR7-10067*).**

**Figure S4: Isolates USA_6 and USA_Ken-4-3 carry independent gene deletions that encompass the *AvrPm8* gene.**

**Figure S5: *AvrPm8* mutations result in gain-of-virulence phenotypes on *Pm8* wheat.**

**Note S1: Splice site mutations**

**Table S1: List of primers used in this study**

**Supplementary Dataset S1: Summary of *B.g. tritici* isolates used in this study**

**Supplementary Dataset S2: List and sequence of gene synthesis constructs used in this study**

