## Supplementary Information for "Doomed by popularity: The broad use of the *Pm8* resistance gene in wheat resulted in hypermutation of the *AvrPm8* gene in the powdery mildew pathogen"

**This PDF file includes:**

Figure S1: Phylogenetic analysis of effector family E003 in *B.g. tritici* isolate ISR_7.

Figure S2: *AvrPm8* (*BgISR7-10067*) is among the top 5% of expressed genes in *B.g. tritici* isolate ISR_7.

Figure S3: Splice site mutations in isolate GBR_JIW2 abolish splicing of *AvrPm8* (*BgISR7-10067*).

Figure S4: Isolates USA_6 and USA_Ken-4-3 carry independent gene deletions that encompass the *AvrPm8* gene.

Figure S5: *AvrPm8* mutations result in gain-of-virulence phenotypes on *Pm8* wheat.

Note S1: Splice site mutations

Table S1: List of primers used in this study

**Other supplementary materials for this manuscript include the following:**

Supplementary Dataset S1: Summary of *B.g. tritici* isolates used in this study

Supplementary Dataset S2: List and sequence of gene synthesis constructs used in this study

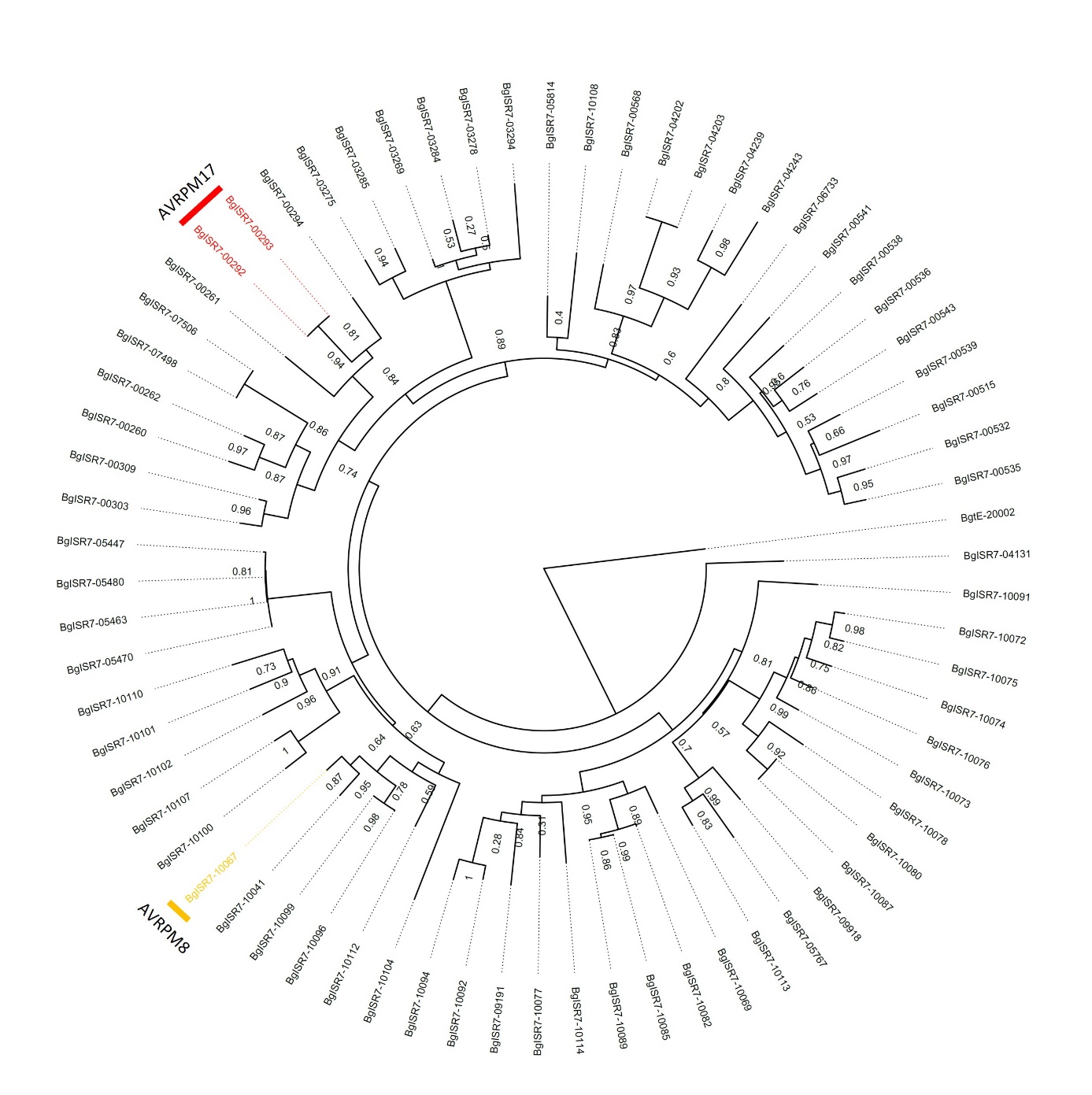

**Figure S1: Phylogenetic analysis of effector family E003 in *B.g. tritici* isolate ISR_7.** Maximum likelihood phylogenetic tree (using the Jones-Taylor-Thorton model) was based on the protein sequences of 69 E003 family members found in ISR_7. The AVRPM3^B2/C2^ (BgtE-20002) protein sequence was used as an outgroup. The Shimodaira-Hasegawa test was used to estimate the local support values to the nearest neighbor interchanges topology and indicated for each branch. AVRPM8 (BgISR7-10067) and AVRPM17 (BgISR7-00292/ BgISR7-00293) homologs in ISR_7 are labelled in yellow and red respectively.

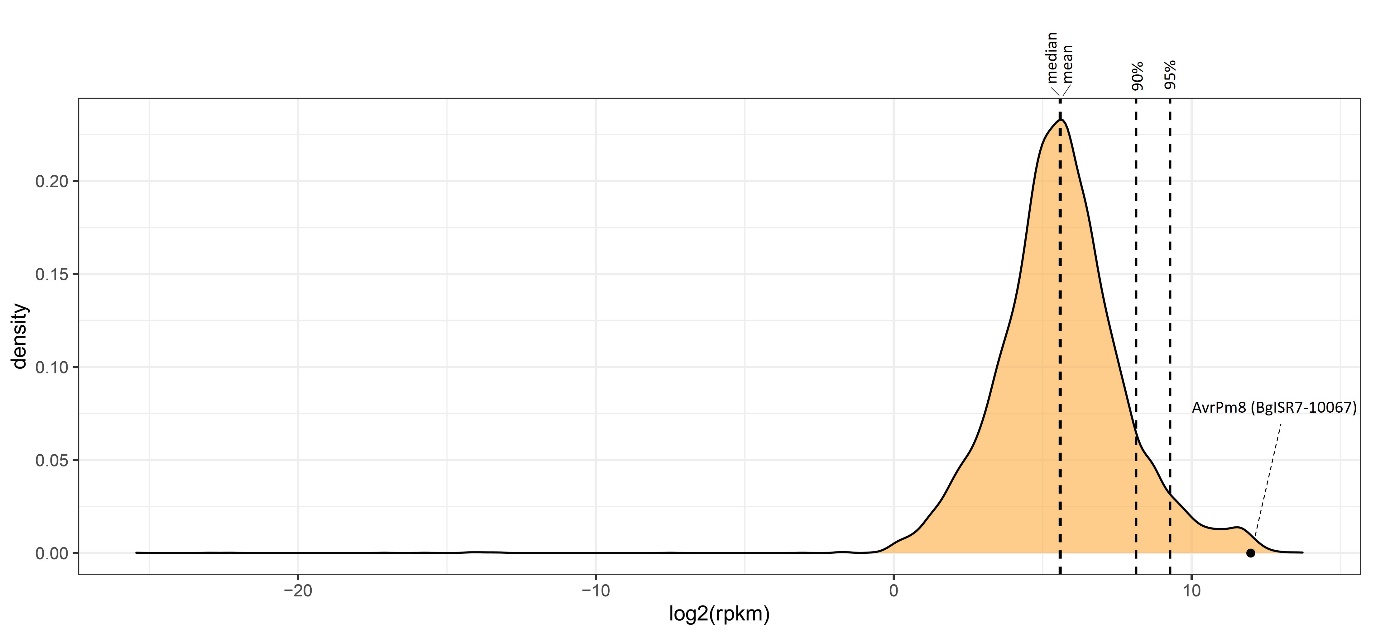

**Figure S2: *AvrPm8 (BgISR7-10067)* is among the top 5% of expressed genes in *B.g. tritici* isolate ISR_7.** Distribution of all expressed genes in ISR_7 (displayed as log2(rpkm) values) at 2 days post infection on wheat cultivar ´Chinese Spring’ are shown. The median, mean, 90% quartile and 95% quartile are indicated by dashed lines. Expression value of *AvrPm8* (*BgISR7-10067*) is indicated by a black dot.

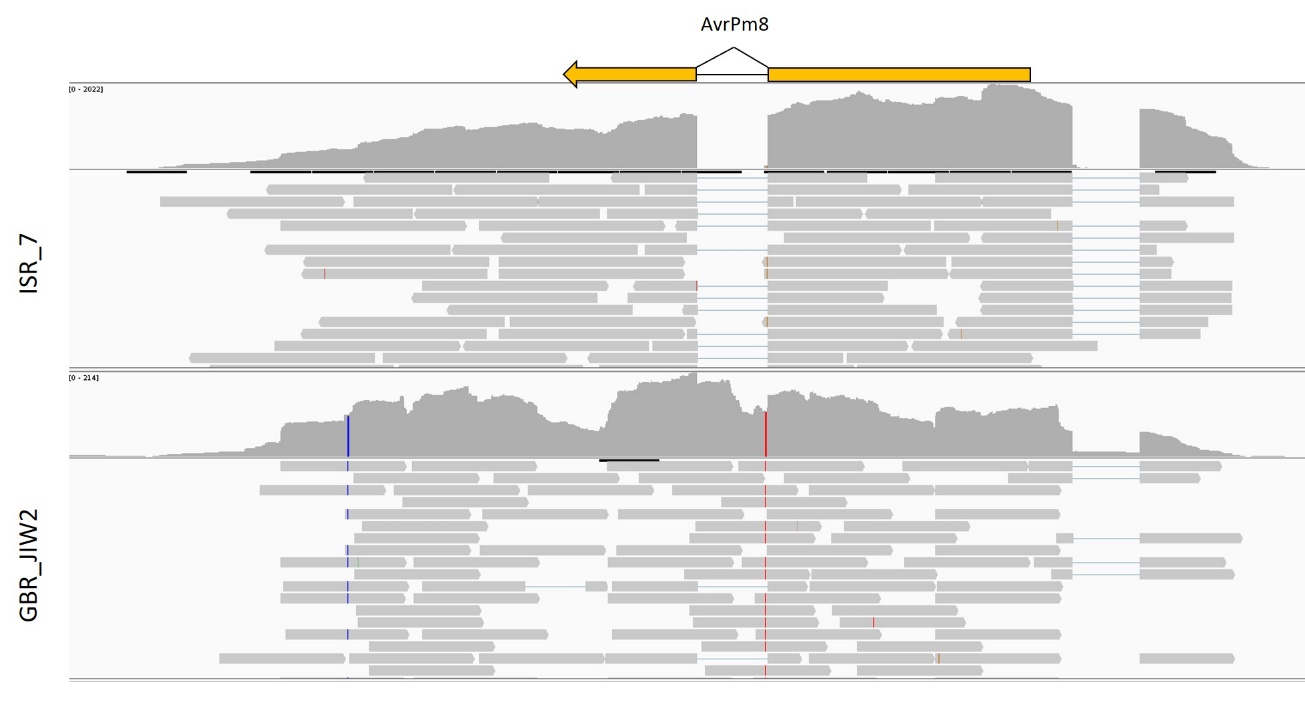

**Figure S3: Splice site mutations in isolate GBR_JIW2 abolish splicing of *AvrPm8* (*BgISR7-10067*).** RNA-sequencing reads of isolates ISR_7 and GBR_JIW2 infecting cultivar “Chinese Spring” at two-day post infection (2dpi) were mapped against the ISR_7 reference genome and visualized with the integrative genome browser (IGV). For each isolate, the top panel depicts coverage of mapped RNA reads, bottom panel depicts individual reads. The gene model of BgISR7-10067 is indicated in yellow above the mapping tracks. Splice site mutation leading to terminal dinucleotides GA-AG (instead of GT-AG in ISR_7) in isolate GBR_JIW2 is visible in red.

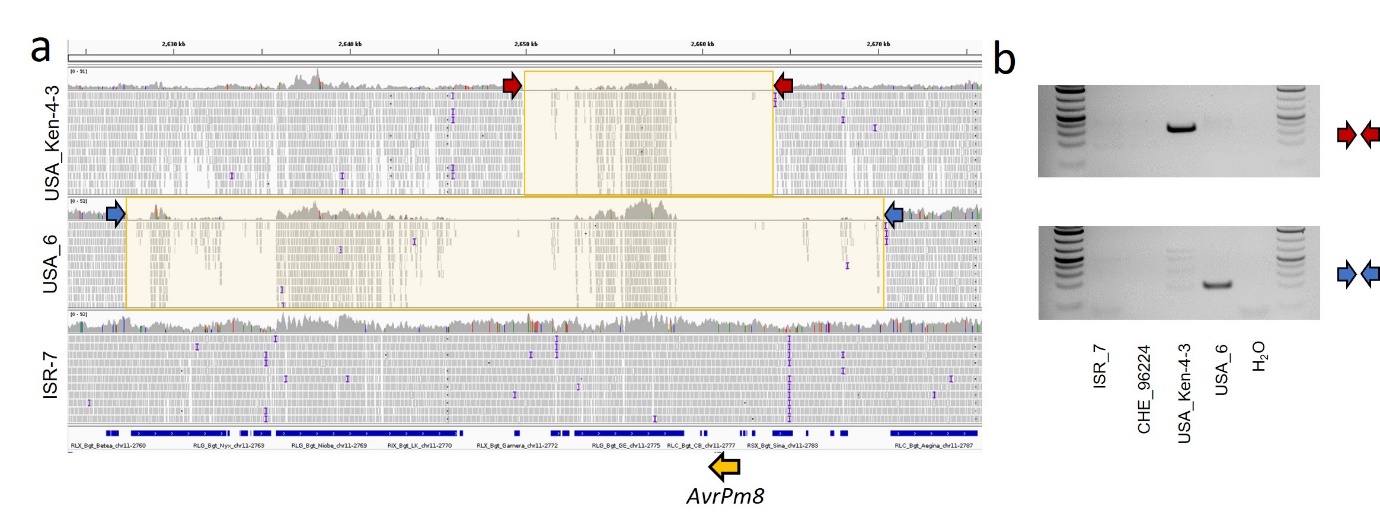

**Figure S4: Isolates USA_6 and USA_Ken-4-3 carry independent gene deletions that encompass the *AvrPm8* gene.** (**a**) Genomic Illumina reads of the isolates USA_Ken-4-3, USA_6 and ISR_7 were aligned against the genome assembly Bgt_genome_v3_16 of isolate CHE_96224 and visualized with the integrative genome browser (IGV). Position of *AvrPm8* is indicated by a yellow arrow. The deleted region for each isolate is indicated by a faint yellow overlay. Transposable element predictions are indicated by blue rectangles below the sequencing tracks. Remaining sequencing reads mapping within the deleted region originate from similar transposable elements elsewhere in the genome. Primers used for PCR validation of the deletion are indicated by red and blue arrows. (**b**) PCR validation on genomic DNA for the deletion events in USA_Ken-4-3 and USA_6 using the specific primer pairs indicated in (a). Absence of deletion specific PCR amplification in ISR_7 and CHE_96224 is shown for comparison.

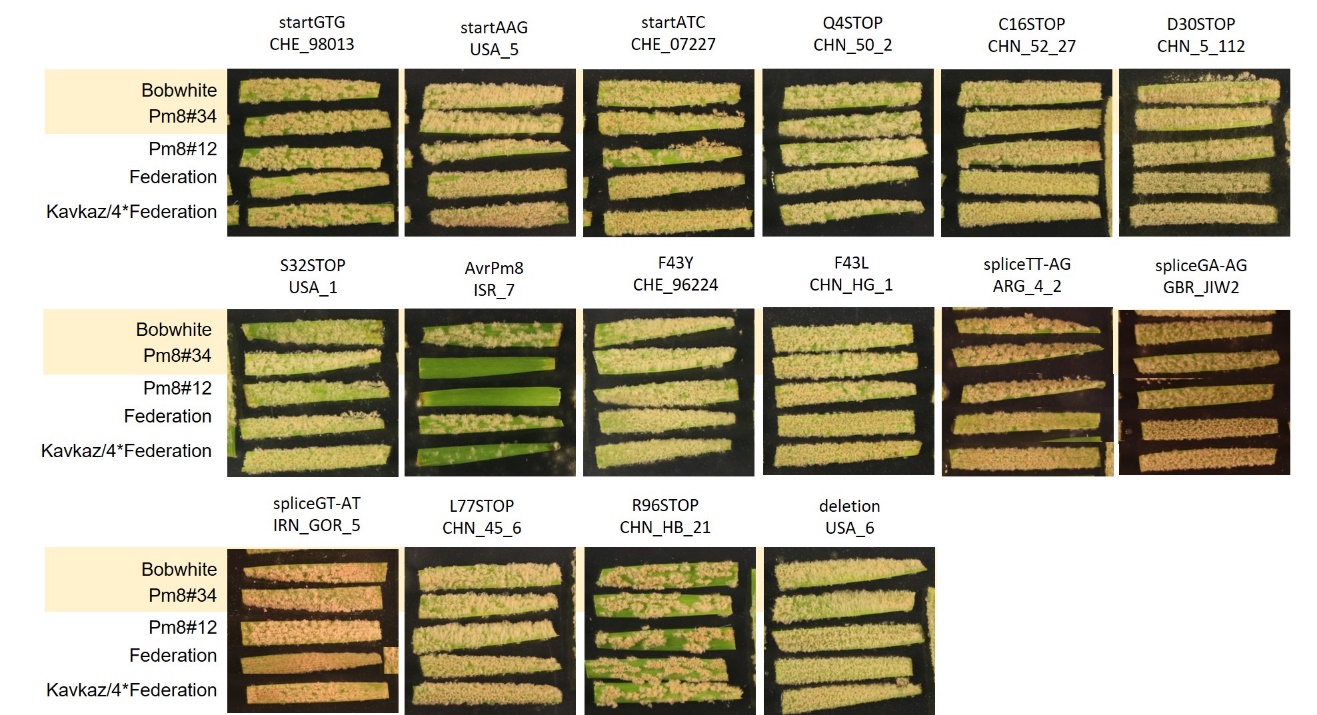

**Figure S5: *AvrPm8* mutations result in gain of virulence phenotypes on *Pm8* wheat.** Virulence phenotype of selected isolates on *Pm8* near-isogenic line ‘Kavkaz/4*Federation’ and two independent *Pm8* transgenic lines ‘Pm8#12’ and ‘Pm8#34’. Susceptible cultivars ‘Federation’ and ‘Bobwhite’ are shown as controls. Sections of the images shown in Figure 4a are highlighted in yellow. Isolate name and corresponding *AvrPm8* mutation are indicated above the picture. Images were taken 8-10 days post infection.
**Note S1: Splice site mutations**

Splicing of pre-mRNA is highly conserved across eukaryotes and relies on the precise recognition of exon-intron junctions by components of the spliceosome complex (Collins & Penny, 2005). Consequently, the sequences at exon-intron boundaries underly strong sequence conservation. In particular the terminal dinucleotides of the intronic sequence canonically consist of GT at the 5’end and AG at the 3’ end in more than 98% of intronic sequences found in eukaryotes (Frey & Pucker, 2020). Mutations in the GT-AG terminal dinucleotides are associated with retention of introns and exon skipping and have been implicated in many hereditary human diseases (Abramowicz & Gos, 2018). Less information is available on the impact of splice site mutations in fungi but given the strong conservation of GT-AG terminal dinucleotides in all eukaryotic kingdoms, consequences are likely similar to the ones observed in animals and plants (Frey & Pucker, 2020). Consistently, a recent study found an EMS-induced splice site mutation in the *AvrSr35* gene of wheat stem rust to cause virulence on *Sr35* wheat lines (Salcedo *et al.*, 2017).

The *AvrPm8* gene contains a single intron of 57 bp with the terminal dinucleotides GT-AG (Fig. 4a). Among the 219 *B.g. tritici* isolates used for *AvrPm8* haplovariant mining, we found 22 isolates with mutations in these highly conserved splice sites, leading to the dinucleotide combinations AT-AG (3 isolates), TT-AG (9), GA-AG (9) and GT-AT (1) while the coding sequence was identical to the recognized *AvrPm8* variant found in ISR_7. We hypothesized, that the disruption of efficient splicing could represent an additional gain of virulence mechanism. Indeed, analysis of RNA sequencing data from GBR_JIW2, carrying the mutated GA-AG splice sites revealed splicing of the intron to be largely abolished (Fig. S3). Ribosomal translation of such unspliced *AvrPm8* mRNA would lead to the encounter of a premature stop codon in the intron and therefore result in a truncated protein. Similar effects on splicing have to assumed for the other identified splice site mutations leading to the terminal dinucleotides AT-AG, TT-AG and GT-AT. Consistent with this hypothesis, isolates carrying splice site mutations consistently exhibited a virulent phenotype on *Pm8* wheat (Fig 4a, Fig. S5).

**Table S1: List of primers used in this study**

| **Name** | **Sequence** | **Purpose** |
| --- | --- | --- |
| LK827 | GCAACCAAACACCTTGAGATGAG | PCR validation of AvrPm8 deletion in USA_Ken-4-3 |
| LK829 | GTGGTGGGATTATTGTGGAGAG | PCR validation of AvrPm8 deletion in USA_Ken-4-3 |
| LK831 | CTCATGAATGAATCATGTCTCTACATGAC | PCR validation of AvrPm8 deletion in USA_6 |
| LK833 | CCTATGATAGTGTTCCGAGCATAG | PCR validation of AvrPm8 deletion in USA_6 |
| LK800 | GAACATCGTATGGATAGCTCCGGCAGGCCTG | SDM primer to create pENTR-Pm8-HA |
| LK801 | CAGATTATGCTTGAAAGGGTGGGCGCGC | SDM primer to create pENTR-Pm8-HA |
